# Deep Sequencing Reveals Compartmentalized HIV-1 in the Semen of Men with and without STI-associated Urethritis

**DOI:** 10.1101/2020.01.28.924225

**Authors:** Olivia D. Council, Shuntai Zhou, Chase D. McCann, Irving Hoffman, Gerald Tegha, Deborah Kamwendo, Mitch Matoga, Sergei L. Kosakovsky Pond, Myron S. Cohen, Ronald Swanstrom

## Abstract

Concurrent sexually transmitted infections (STI) can increase the probability of HIV-1 transmission primarily by increasing the viral load present in semen. In this study, we explored the relationship of HIV-1 in blood and seminal plasma in the presence and absence of urethritis and after treatment of the concurrent STI. Primer ID deep sequencing of the V1/V3 region of the HIV-1 *env* gene was done for paired blood and semen samples from ART-naïve men living in Malawi with (n = 19) and without (n = 5) STI-associated urethritis; for a subset of samples full length *env* genes were generated for sequence analysis and to test entry phenotype. Cytokine concentrations in the blood and semen were also measured, and a reduction in the levels of pro-inflammatory cytokines was observed following STI treatment. We observed no difference in the prevalence of diverse compartmentalized semen-derived lineages in men with or without STI-associated urethritis, and these viral populations were largely stable during STI treatment. Clonal amplification of one or a few viral sequences accounted for nearly 50% of the viral population indicating a recent bottleneck followed by limited viral replication. We documented a case of superinfection where the new strain was restricted to the genital tract. We conclude that the male genital tract is a site where virus can be brought in from the blood, where localized sustained replication can occur, where a superinfecting strain can persist, and where specific genotypes can be amplified perhaps initially by cellular proliferation but further by limited viral replication.

**Importance:** HIV-1 is a sexually transmitted infection that co-exists with other STIs. Here we examine the impact of a concurrent STI resulting in urethritis on the HIV-1 population within the male genital tract. We found that viral populations remain largely stable even with treatment of the STI. These results show that viral populations within the male genital tract are defined by factors beyond transient inflammation associated with a concurrent STI.

## Introduction

Nearly two million new HIV-1 infections occur worldwide every year, predominately through sexual transmission (1). Therefore, understanding the genotypic and phenotypic properties of HIV-1 present in the male genital tract is vital for treatment and prevention strategies. It has been well-established that the probability of sexual transmission of HIV-1 increases with an increasing viral load (2–5), and there are several factors that can influence the concentration of viral RNA present in semen. For example, stage of disease (6), CD4+ T cell count (7), and the presence of inflammatory conditions (such as concurrent sexually transmitted infections [STI]) have all been demonstrated to increase the semen viral load (reviewed in (8)).

For semen-mediated transmission events, the transmitted/founder virus is most proximal to the male genital tract at the time of transmission. Thus, the origin of virus in the male genital tract is relevant to a fuller understanding of HIV-1 transmission. Often the virus present in semen is similar to virus found in the blood (an equilibrated population), but there is also evidence that the male genital tract is able to support independent replication of HIV-1. This fact is inferred from observations of genetically distinct, or compartmentalized, HIV-1 populations in semen, as compared to the virus found in the blood and other anatomical compartments (9–15). In addition, several studies (16, 17) have reported the presence of HIV-1 RNA in the semen of men on suppressive antiretroviral therapy, with undetectable blood plasma viral loads, implying that the male genital tract can influence viral replication independent of the periphery and harbor an independent viral reservoir. It is therefore important to elucidate the factors that promote the establishment and maintenance of compartmentalized viral lineages in the male genital tract.

In the current study, we examined the effects of STI-associated urethritis on the establishment and maintenance of compartmentalized lineages in the male genital tract by comparing viral sequences in the blood and in seminal plasma using deep sequencing technology with Primer ID (18, 19). We explored the possibility that STI-associated inflammation could act to recruit CD4+ T cells into the genital tract, thereby promoting a mixing of viral populations in the blood and semen with a concomitant reduction in apparent compartmentalization, or conversely the influx of cells could enhance the replication of locally produced virus and increase compartmentalization. We also examined the viral population dynamics between blood and semen over time to determine whether antibiotic treatment of the concurrent STI would impact HIV-1 compartmentalization. We detected no difference between the proportions of men who had compartmentalized, semen-derived lineages, grouped by the presence or absence of urethritis. Furthermore, antibiotic treatment of the STI did not observably impact the population dynamics between the blood and the semen, at least in the short term. We conclude that STI-associated inflammation is not a driving factor behind the establishment or maintenance of compartmentalized lineages in the semen and that independent viral replication can occur independently of inflammatory conditions.

## Methods

### Ethics Statement and Source of Clinical Samples

Blood and semen samples were collected as part of a study examining the effects of genital tract inflammation on HIV-1 semen viral load (20). The study was approved by the Institutional Review Board at the University of North Carolina at Chapel Hill. A subset of STI samples (12/19) were previously examined via a heteroduplex tracking assay (9), and 2/5 control samples were previously examined via single genome amplification (SGA) (10).

### Deep Sequencing with Primer ID

Deep sequencing with Primer ID was performed as previously described (19). Briefly, viral RNA was extracted from seminal and blood plasma using the QIAamp Viral RNA Extraction Kit (Qiagen). Based on viral loads, up to 5,000 RNA copies (range: 196-5,000, mean: 3,161) were used for cDNA synthesis. cDNA was synthesized using the *env* V1/V3 Primer ID primer (HXB2 positions 6585-7208): 5’-GTGACTGGAGTTCAGACGTGTGCTCTTCCGATCTNNNNNNNNNCAGTCCATTTTGCTCTACTAATGTTACAATGTGC-3’ and SuperScript III Reverse Transcriptase (Invitrogen). The final cDNA reaction contained the following: 0.5 mM dNTP mix (KAPA) 0.25 μM V1-V3 reverse primer, 5 mM DTT, 6 U RNaseOUT, and 30 U SuperScript III RT in a total volume of 60 μl. Initially a mixture containing dNTPs, cDNA primer and RNA template was incubated at 65°C for 5 minutes, followed by 4°C for 2 minutes. Then DTT, RNaseOUT and SuperScript III were added and the reactions were incubated for one hour at 50°C, followed by one hour at 55°C. Samples were then heated to 70°C for 15 minutes to inactivate the SuperScript III prior to addition of RNase H (2 units) and a final incubation at 37°C for 20 minutes. cDNA was purified using Agencourt RNAclean XP beads (Beckman Coulter) at a volume ratio of 0.6:1 beads: cDNA. The beads were washed four times with 70% ethanol. Purified cDNA was eluted in 24 μl mοlecular grade water (Corning), and the purification was repeated with a bead:cDNA ratio of 0.6:1. The purified cDNA was again eluted in 24 μl molecular grade water and stored at −20°C. All of the cDNA (24 μl) was used for PCR amplification. KAPA 2G Robust HotStart Polymerase was used as the first-round PCR enzyme along with the following forward primer: 5’-GCCTCCCTCGCGCCATCAGAGATGTGTATAAGAGACAGNNNNTTATGGGATCAAAGCCTAAAGCCATGTGTA-3’ corresponding to the HIV-1 *env* V1/V3 region. Following amplification, PCR products were purified using AmpureXP beads (Beckman Coulter) at a ratio of 0.7:1 beads: DNA. Beads were washed four times using 70% ethanol, and the purified DNA was eluted in 50 μl of DNase-free water (Corning). The second round of PCR consisted of 2 μl of purified first-round PCR product along with the KAPA HiFi Robust Polymerase enzyme and served to incorporate MiSeq adaptors and index oligonucleotides that allowed for multiplexing of samples.

### MiSeq Library Preparation and Quality Control

Amplicons were visualized on a 1.2% agarose gel. Gel extraction was performed using the MinElute Gel Extraction Kit (Qiagen) according to manufacturer’s instructions. Purified DNA was eluted in 10 μl of EB Buffer (Qiagen) and quantified using the Qubit dsDNA Broad Range Assay (Thermo Fisher). Samples were pooled in equimolar concentrations and the final library was purified using AmpureXP beads at a ratio of 0.7:1 beads: DNA. Libraries were submitted to the UNC High Throughput Sequencing Facility for generation of 2×300 base paired-end reads using the Illumina MiSeq platform.

### Phylogenetic and Compartmentalization Analyses

Compartmentalization of viral populations was assessed using two tree-based methods: the Slatkin-Maddison (S-M) test (21) and the presence of a genetically diverse, semen-derived lineage. The S-M test was performed on phylogenetic trees that had equal numbers of semen and blood-derived V1/V3 sequences, after collapsing identical sequences in each compartment to focus on diverse populations rather than clonally amplified populations. The standard Slatkin-Maddison test was modified to account for the structure of the tree, with the leaves of each node being permutated sequentially before inferring migrations (Pond et al, manuscript in preparation, https://github.com/veg/hyphy-analyses/tree/master/SlatkinMaddison). Trees were considered compartmentalized if 10,000 permutations of the Standard Slatkin-Maddison test or 50,000 permutations the Structured Slatkin-Maddison test yielded a p-value <0.05 and there was a semen-derived, genetically diverse lineage. Both S-M tests are implemented in the standard analysis “sm” in HyPhy v2.5.

### Single Genome Amplification

Single genome amplification (SGA, or template end-point dilution PCR) was performed as previously described (10). Briefly, viral RNA was extracted using a QIAamp viral RNA extraction kit (Qiagen). cDNA was synthesized using an oligo(dT) primer and SuperScript III RT (Invitrogen). Template cDNA was diluted such that <30% of reactions were positive in the subsequent PCR. Nested PCR was performed using Platinum *Taq* High Fidelity polymerase (Invitrogen) and the following primers: PCR-1: 5’-GGGTTTATTACAGGGACAGCAGAG-3’ (Vif1) and 5’-TAAGCCTCAATAAAGCTTGCCTTGAGTGC-3’ (OFM19), PCR-2: 5’-GGCTTAGGCATCTCCTATGGCAGGAAGAA-3’ (EnvA) and 5’-ACACAAGGCTACTTCCCTGGATTGGCAG-3’ (EnvN). SGA products were fully sequenced from both directions to confirm the presence of a single template. Amplicons with evidence of multiple templates (i.e., double peaks on the chromatogram) were not used in downstream applications.

### Construction of HIV-1 *env* clones

Amplicons of the full-length HIV-1 *env* gene from the first round PCR with confirmed sequences were subjected to an additional round of PCR using the Phusion hot-start high fidelity DNA polymerase (Invitrogen) and the primers cEnvA (5’-CACCGGCTTAGGCATCTCCTATACCAGGAAGAA-3’) and EnvN (5’-CTGCCAATCAGGGAAGTAGCCTTGTGT-3’) following the manufacturer’s instructions. HIV-1 *env* amplicons were then gel purified using the Qiagen QIAQuick Gel Extraction Kit. An aliquot of 50 ng of purified HIV-1 *env* DNA was used to clone into the pcDNA 3.1D/V5-His-TOPO vector (Invitrogen) and MAX Efficiency Stlb2 competent cells (Life Technology) per the manufacturer’s instructions.

### Env-pseudotyped viruses

Env*-*pseudotyped luciferase reporter viruses were generated by co-transfection of 810 ng of an *env* expression vector and 810 ng of pZM247Fv2Δenv backbone (22) using 293T cells and the Fugene 6 reagent and protocol (Promega). Five hours after transfection, the medium was changed. Forty-eight hours after transfection, the medium was harvested, filtered through a 0.45 μm filter, and aliquoted into 0.6 ml tubes. Aliquots were stored at −80°C until use.

### Single-cycle infection of 293-Affinofile cells

The ability of HIV-1 Env proteins to mediate infection of cells expressing low densities of CD4 was assessed as previously described (23–25). Briefly, experiments were carried out in black, flat-bottomed, 96-well plates. A solution of 100 μl of 293-Affinofile cells at a density of 1.8×10^5^ cells/ml was added to the inner 60 wells of each 96-well plate. All 293-Affinofile cells were induced to express high levels of CCR5 expression using Ponesterone A. CD4 expression was induced in half of the cells using Doxycycline. Twenty-four hours after CCR5 and/or CD4 induction, cells were spinoculated (26) with previously titered Env-pseudotyped viruses (849 x g for 2 hours at 37°C). Following spinoculation, cells were incubated at 37°C for forty-eight hours. Cells were then washed twice with PBS and lysed with 1x Renilla luciferase assay lysis buffer diluted in distilled water. Following lysis, plates were kept at −80°C overnight. The following day, plates were thawed at room temperature and read using a luminometer. A 50 μl aliquot of Renilla assay reagent was injected into the luminometer per well, and relative light units (RLUs) were recorded over 5 seconds with a 2-second delay.

### Cytokine Evaluation

Cytokine concentrations in blood plasma and seminal plasma were quantified using a Luminex® bead-based multiplex assay (R&D Systems). Specifically, TNF-α, IL-6, CXCL10, IL-10, CCL2, IL-1β, and IFN-γ concentrations were determined. All assays were performed following the manufacturer’s instructions.

### Data availability

The full length *env* gene sequences will be deposited in GenBank on acceptance and the accession numbers included in proof. The MiSeq sequences will be deposited in the Sequencing Read Archive and the accession numbers included in proof.

## Results

### Participant characteristics and sequence generation

Participants were part of a cohort of men based in Malawi that was established to examine the effect of STI-associated urethritis on seminal plasma HIV-1 viral load (20). In order to examine the relationship between urethritis associated with a concurrent sexually transmitted infection (STI) and the presence of compartmentalized virus in the genital tract, we selected a subset of men with (n = 19) and without (n = 5) STI-associated urethritis, with the sample size determined by availability of sufficient seminal plasma. All participants were chronically infected with HIV-1, and antiretroviral therapy (ART) naive, as ART was not available in Malawi at the time of the study.

Participant characteristics are shown in Table 1. There was no difference in the blood viral load, semen viral load, or CD4+ T cell count between the two groups at baseline. HIV-1 RNA was extracted from paired blood plasma and seminal plasma, and Illumina MiSeq deep sequencing with Primer ID was used to generate HIV-1 *env* V1/V3 amplicons. The deep sequencing output was collapsed into Template Consensus Sequences (TCS) for each Primer ID recovered to create a highly accurate sequence for each original RNA template sampled. An average of 62 TCSs were obtained from each compartment (blood and semen) for each participant (range: 12-200), giving us 95% power to detect minor populations present in most samples at a 1.5-5% frequency.

**Table 1.** Relevant clinical information for the particpants analyzed. ^a^Cells per microliter of blood. ^b^Diagnosed STI. Gon, gonorrhea; Tri, trichomonas; Ulc, genital ulcers; NPI, no pathogen identified. ND, not done.

|  | Before STI Treatment |  | After STI Treatment |  | CD4 Count<sup>a</sup> | Diagnosed STI<sup>b</sup> |
| --- | --- | --- | --- | --- | --- | --- |
|  | Plasma Viral Load<br>(log<sub>10</sub> copies/mL) | Semen Viral Load<br>(log<sub>10</sub> copies/mL) | Plasma Viral Load<br>(log<sub>10</sub> copies/mL) | Semen Viral Load<br>(copies/mL) |  |  |
| **Urethritis (n = 19)** |  |  |  |  |  |  |
| S003 | 5.23 | 5.48 | 5.51 | 4.8 | ND | Gon., Ulc |
| S018 | 5.59 | 4.70 | 5.40 | 4.96 | 476 | Gon, Tri, Ulc |
| S019 | 4.18 | 5.48 | 4.34 | 4.32 | 712 | Gon |
| S031 | 4.95 | 4.62 | 4.63 | 4.92 | 333 | Gon, Ulc |
| S053 | 6.11 | 6.08 | 5.88 | 5.38 | 318 | Tri, Ulc |
| S070 | 4.73 | 3.92 | 4.96 | 4.46 | 670 | Gon |
| S073 | 4.59 | 4.60 | 4.51 | 4.23 | 275 | Gon |
| S075 | 5.59 | 5.20 | 5.90 | 4.08 | ND | NPI |
| S101 | 5.83 | 5.89 | 5.79 | 5.20 | 235 | Gon |
| S103 | 3.95 | 5.00 | 5.11 | 4.79 | 454 | Gon |
| S146 | 4.79 | 4.96 | 5.58 | 5.53 | 258 | Ulc |
| S172 | 4.80 | 5.18 | 4.28 | 4.62 | 344 | Gon |
| S029 | 5.23 | 4.71 | ND | ND | 496 | Gon |
| S017 | 5.79 | 4.40 | ND | ND | 178 | NPI |
| S039 | 5.56 | 5.04 | ND | ND | ND | Gon |
| S047 | 5.77 | 4.92 | ND | ND | 469 | Gon |
| S099 | 5.57 | 5.82 | ND | ND | ND | Gon, Tri |
| S148 | 5.08 | 5.98 | ND | ND |  | Gon |
| S067 | 4.90 | 5.04 | ND | ND |  | Gon |
| **No Urethritis (n =5)** |  |  |  |  |  |  |
| C019 | 7.00 | 5.72 | 5.72 | 6.76 | 599 | None |
| C073 | 4.82 | 3.76 | ND | ND | ND | None |
| C082 | 5.49 | 4.20 | ND | ND | 305 | None |
| C111 | 4.84 | 5.95 | ND | ND | 210 | None |
| C061 | 5.56 | 6.51 | ND | ND | 262 | None |

### Compartmentalized, semen-derived lineages are observed in men with and without urethritis

As we were primarily interested in identifying diverse compartmentalized lineages, which represent independent replication over a period of time, rather than compartmentalized lineages that consist primarily of clonal sequences, we initially collapsed sequences that were identical to within one nucleotide into a single haplotype. After identical sequences were collapsed, an equal number of blood-derived and semen-derived sequences were used to construct neighbor-joining phylogenetic trees for each participant, allowing us to compare the two populations at equivalent sampling depth. Compartmentalization was assessed using both the Slatkin-Maddison test (21), and the Structured Slatkin-Maddison test (Pond et al, in preparation, https://github.com/veg/hyphy-analyses/tree/master/SlatkinMaddison), which has been modified to reduce potentially spurious compartmentalization detection in trees with large numbers of sequences. When both tests resulted in a P value < 0.05, the tree was deemed compartmentalized. When one test indicated compartmentalization while the other did not, trees were inspected visually for the presence of diverse, semen-dominated lineages.

Among the 24 men, we observed varying degrees of compartmentalization, ranging from near-complete separation of blood and semen-derived sequences, to minor compartmentalization in 6/24 (25%) participants. In men with urethritis, compartmentalization was detected in 5/19 (26%) men, while viral populations were equilibrated between the blood and semen in 13/19 (68%) men. One individual with urethritis was superinfected, with the superinfecting population constituting a distinct, semen-only lineage. In men without urethritis, we observed minor compartmentalization in 1/5 (20%) individuals, and equilibrated viral populations in 4/5 (80%) individuals. Thus, both compartmentalized and equilibrated HIV-1 populations were found in men with and without urethritis (Figure 1 and Table 2) at statistically indistinguishable frequencies, although the generalizability of this conclusion is limited by the number of samples studied.

**Figure 1.**
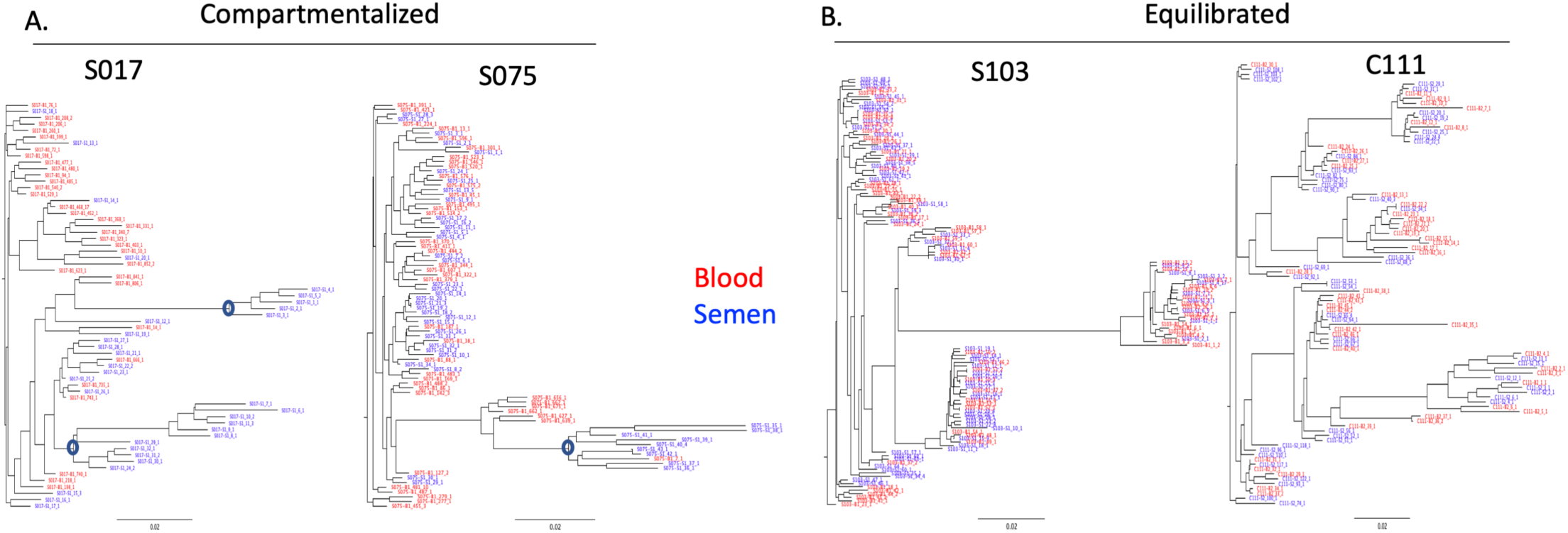
Representative neighbor-joining *env* V1/V3 phylogenetic trees depicting compartmentalization between the blood and semen-derived lineages (A) and equilibration between blood and semen-derived lineages (B). Blood-derived sequences are shown in red, semen-derived sequences are shown in blue. Circles indicate compartmentalized nodes.

**Table 2.** Summary of the methods used to determine the relationship between blood-derived and semen-derived HIV-1 *env* V1/V3 sequences. Bolded values indicate statistically significant compartmentalization as determined by the indicated method.

|  | Before STI Treatment |  |  |  | After STI Treatment |  |  |  |
| --- | --- | --- | --- | --- | --- | --- | --- | --- |
|  | # HIV *env*<br>V1/V3<br>sequences | Structured<br>Slatkin-Maddison<br>value | Standard<br>Slatkin-Maddison<br>value | Visual Inspection | # HIV *env*<br>V1/V3<br>sequences | Structured<br>Slatkin-Maddison<br>value | Standard<br>Slatkin-Maddison<br>value | Visual Inspection |
| **Urethritis**<br>**(n = 19)** |  |  |  |  |  |  |  |  |
| S003 | 187 | 0.15 | **0.0009** | Equilibrated | 25 | 0.788 | 0.392 | Equilibrated |
| S018 | 120 | 0.327 | 0.186 | Equilibrated | 24 | 0.926 | 0.819 | Equilibrated |
| S019 | 48 | 0.076 | **0.004** | Equilibrated | 22 | 0.335 | **0.03** | Equilibrated |
| S031 | 200 | **0** | **0** | Superinfection | 306 | **0** | **0** | Superinfection |
| S053 | 29 | 0.461 | **0.027** | Equilibrated | 23 | 0.396 | 0.111 | Equilibrated |
| S070 | 15 | 0.946 | 0.539 | Minor compartment. | 12 | 0.262 | **0.017** | Minor Compartment. |
| S073 | 24 | 0.882 | 0.734 | Equilibrated | 8 | 0.428 | 0.428 | Equilibrated |
| S075 | 43 | 0.263 | **0.003** | Minor compartment. | 42 | 0.515 | **0.06** | Minor Compartment. |
| S101 | 13 | 0.653 | 0.15 | Equilibrated | 103 | **0** | **0** | Compartmentalized |
| S103 | 62 | 0.904 | 0.902 | Equilibrated | 10 | 0.322 | 0.098 | Equilibrated |
| S146 | 71 | 0.251 | **0.004** | Minor compartment. | 45 | 0.36 | **0.033** | Minor Compartment. |
| S172 | 32 | 0.805 | 0.277 | Equilibrated | 24 | 0.96 | 0.89 | Equilibrated |
| S029 | 47 | 0.713 | 0.17 | Equilibrated | ND | ND | ND | ND |
| S017 | 32 | **0.003** | **0** | Minor compartment. | ND | ND | ND | ND |
| S039 | 19 | 0.825 | 0.61 | Equilibrated | ND | ND | ND | ND |
| S047 | 25 | 0.916 | 0.383 | Equilibrated | ND | ND | ND | ND |
| S099 | 200 | **0.002** | **0.0009** | Compartmentalized | ND | ND | ND | ND |
| S148 | 82 | 0.294 | **0.014** | Equilibrated | ND | ND | ND | ND |
| S067 | 98 | 0.203 | **0.01** | Equilibrated | ND | ND | ND | ND |
| **No**<br>**Urethritis**<br>**(n = 5)** |  |  |  |  |  |  |  |  |
| C019 | 25 | 0.787 | 0.369 | Equilibrated | 32 | 0.649 | **0.029** | Equilibrated |
| C073 | 15 | 0.795 | 0.111 | Equilibrated | ND | ND | ND | ND |
| C082 | 12 | 0.537 | 0.248 | Equilibrated | ND | ND | ND | ND |
| C111 | 46 | 0.572 | **0.025** | Equilibrated | ND | ND | ND | ND |
| C061 | 34 | 0.404 | **0.006** | Minor Compartment. | ND | ND | ND | ND |

### HIV-1 population dynamics between blood and semen remain largely stable after STI treatment

To examine the effects of antibiotic treatment of the STI on HIV-1 population dynamics, we compared pre-treatment and post-treatment time points in 13 men (12 with urethritis, 1 without urethritis). Samples were obtained an average of 12 days after antibiotic treatment had been initiated (range: 7-14 days). Neighbor-joining phylogenetic trees were built as described above, and the relationship between blood and semen-derived sequences (i.e., equilibrated or compartmentalized) was determined at each time point. In 12/13 men, the relationship did not change following STI treatment (Figure 2). In one individual, S101, semen and blood-derived lineages were equilibrated in the pre-STI treatment time point, but compartmentalized post-treatment, as determined by both the Standard Slatkin-Maddison test and the Structured Slatkin-Maddison test (p < 0.0001, Table 2).

**Figure 2.**
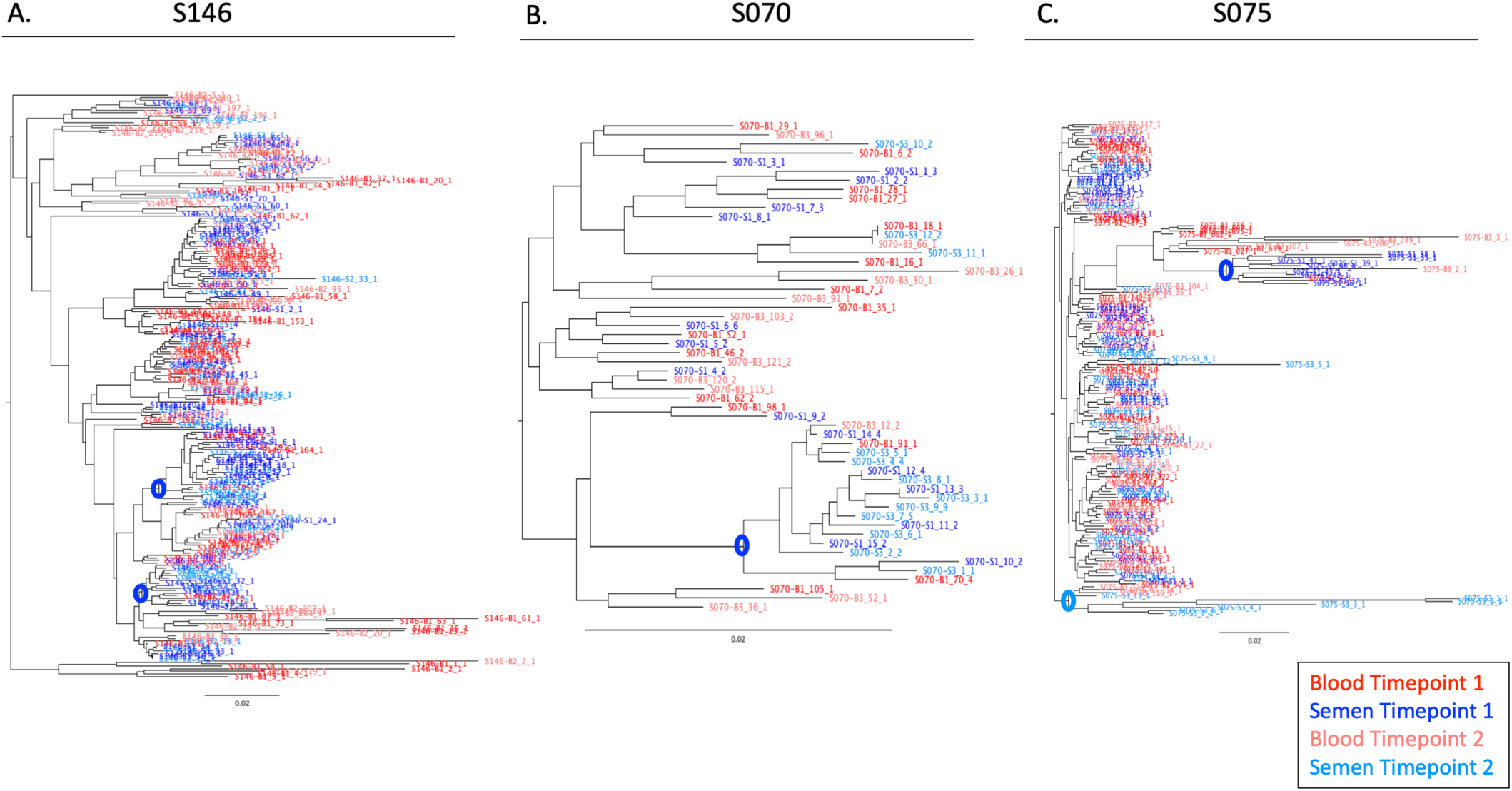
HIV-1 population dynamics between blood and semen remain unchanged after STI treatment. Neighbor-joining phylogenetic trees depicting blood-derived *env* V1/V3 sequences (shades of red) and semen-derived *env* V1V3 sequences (shades of blue). Sequences from before and after STI treatment are shown. In (A) and (B), the same compartmentalized lineage appears in the semen before and after STI treatment (circled nodes). In (C), a different semen-derived, compartmentalized lineage is observed in the pre and post STI-treatment time points.

Next, we compared within-compartment viral diversity before and after STI treatment. To this end, we inferred neighbor-joining phylogenetic trees containing equal numbers of semen-derived sequences from the pre and post STI treatment time points for the 13 men described above. Both the Standard Slatkin-Maddison and the Structured Slatkin-Maddison tests were performed on the trees in order to determine whether the pre- and post-STI treatment semen sequences constituted distinct clades. In 12 of 13 men, the semen populations before and after STI treatment were not significantly different from one another – i.e., populations that existed before STI treatment were still readily observable after STI treatment. Of particular interest were the individuals with compartmentalized, semen-derived lineages. In 2 of the 3 men with compartmentalized lineages at both time points, the lineage that was responsible for the compartmentalization was the same before and after STI treatment (Figure 2A and 2B). Thus, not only was the relationship between compartments unchanged, but the specific lineages themselves persisted. However, in one individual the compartmentalized, semen-derived lineage that was detected before STI treatment was not detected at the second time point but a new compartmentalized lineage was observed (Figure 2C).

### Clonal amplification of blood and semen-derived sequences is observed in men with and without urethritis

As we were primarily interested in the presence of diverse, compartmentalized lineages, rather than compartmentalized lineages comprised of a clonally expanded population, we collapsed sequences that were identical to within one nucleotide into a single haplotype. In doing so, we observed that a large proportion of both blood and semen-derived V1/V3 sequences were identical or nearly identical. Such an observation could be made because of the PCR amplification step prior to sequencing where the original templates are repetitively sequenced, a phenomenon called PCR resampling; however, the use of Primer ID to tag each original templates before PCR avoids this problem allowing us to infer the presence of identical or near identical sequences within the viral population in vivo. For blood-derived sequences, a mean of only 41% and 48% of sequences were unique in men with and without urethritis, respectively (p = 0.4237, Figure 3A). For semen-derived sequences, a mean of only 44% and 62% of sequences were unique in men with and without urethritis, respectively (p = 0.086, Figure 3C). The proportion of unique sequences observed in blood and semen remained stable before and after STI treatment in men with urethritis (Figure 3B and 3D). This result indicates that a significant fraction of the population in each compartment was in a genetic bottleneck or had recently gone through a bottleneck.

**Figure 3.**
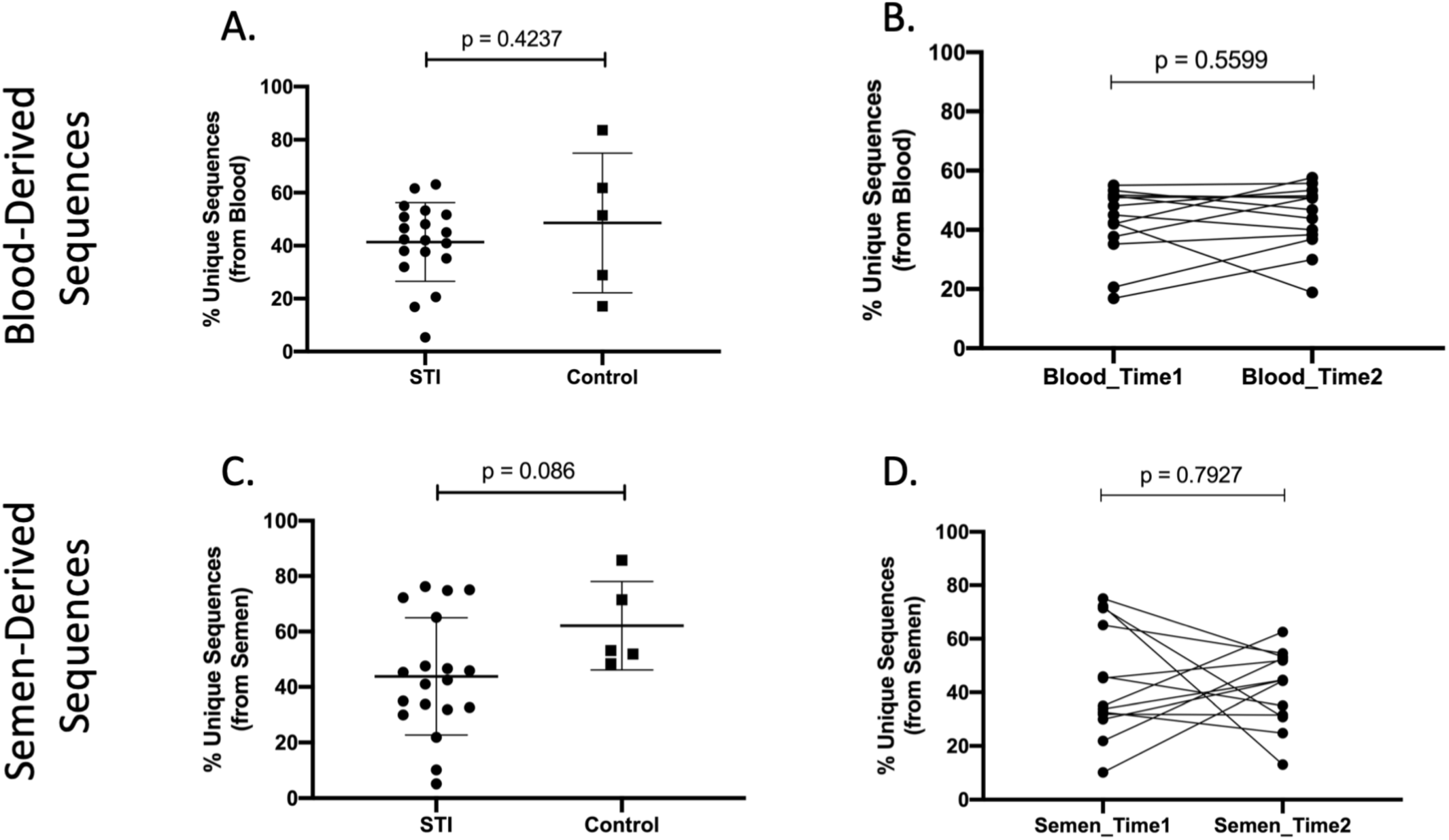
Clonal amplification of identical sequences is observed in both blood and semen-derived viruses in men with and without urethritis. An analysis of the percent of V1/V3 sequences that are not identical, derived from blood (A), or semen (C). The percent of unique sequences remains stable in overtime in both the blood (B) and semen (D). An unpaired t test was used to generate the indicated p values.

We considered the possibility that the short *env* V1/V3 amplicon (527 bases) would over-estimate the percentage of sequences that were identical across the entirety of *env*. To evaluate this possibility, we performed single genome amplification (SGA) of full-length HIV-1 *env* genes (∼2500 bases) from the blood and semen of four men (3 with urethritis and 1 without). We obtained an average of 30 full-length *env* sequences from each participant. In two of the four cases, we observed identical sequences. When we trimmed the full-length sequences and analyzed only the V1/V3 region used in our deep sequencing, we observed identical or nearly identical sequences in all four participants. Sequences that were identical in the V1/V3 region but different across the entire envelope had only a few nucleotide changes between them, consistent with the low-level diversity generated from recent viral replication from a unique ancestor/bottleneck (Figure 4, and Supplemental Figures 1-3). Thus, while examining only the V1/V3 region does increase the number of sequences that appear identical, the overall viral diversity of those variants is low and consistent with recent clonal expansion involving a bottleneck with subsequent viral replication to introduce modest diversity. In a control experiment we generated 8 *env* amplicons from virus produced from the cell line 8E5, which contains a single defective viral genome. When the 8 amplicons were sequenced we observed a single substitution mutation and a single frameshift mutation (data not shown). The low-level diversity observed in the viral populations in vivo were in most cases greater than the level observed in the control amplification, consistent with ongoing viral replication after a recent bottleneck rather than just virus production from a clonally expanded cell.

**Figure 4.**
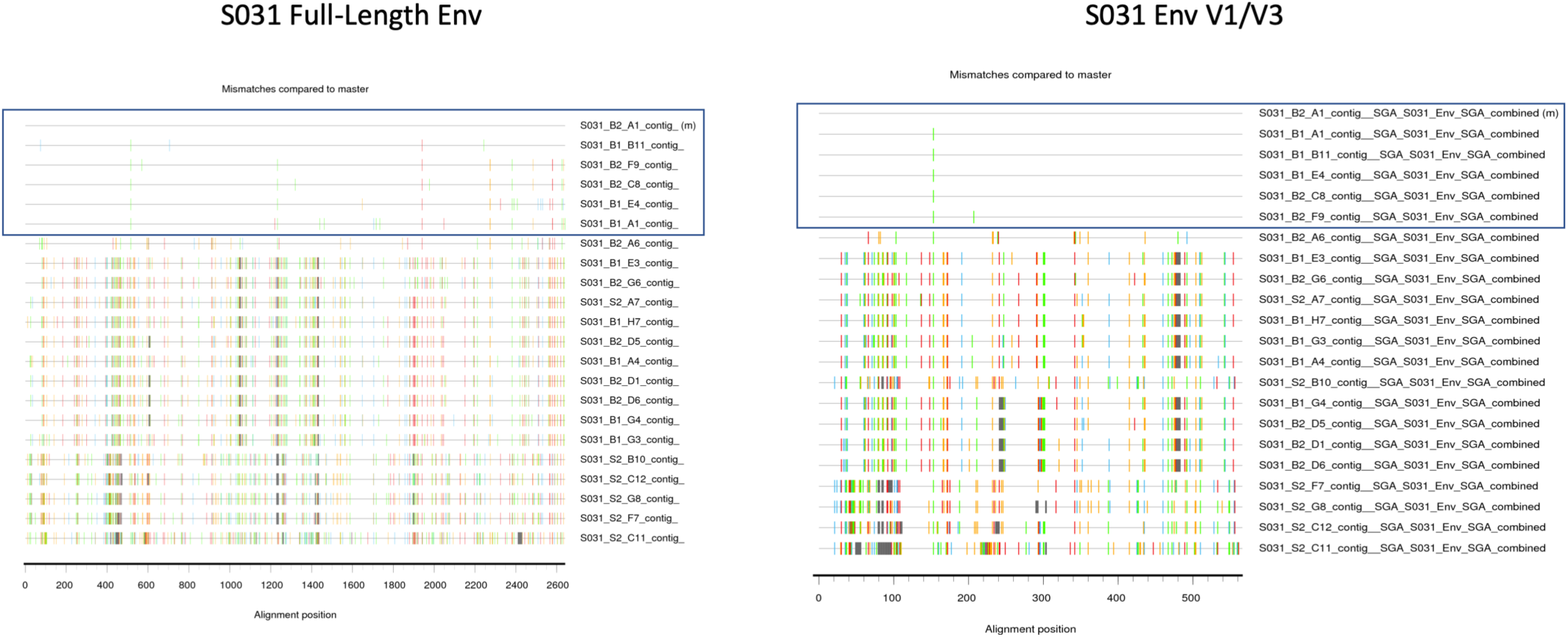
Highlighter plot of paired full-length *env* and *env* V1/V3 sequences. SGA-derived full-length envelope sequences and the corresponding V1/V3 region only are shown on the left and right, respectively. Boxed sequences represent those that are identical in the V1/V3 amplicon, and nearly identical over the full envelope amplicon.

### Semen-derived HIV-1 envelopes are T-cell tropic

HIV-1 primarily infects CD4+ T cells, which have a high density of the CD4 protein on their cell surface that is typically required by the virus for efficient entry. However, viruses that have been replicating independently in anatomically distinct regions such as the central nervous system where CD4+ T cells are less abundant, can evolve the ability to enter cells expressing lower densities of CD4, such as macrophages. This has been observed for compartmentalized lineages derived from both the CNS (27) and, in one case, the male genital tract (28). In order to determine whether compartmentalized, semen-derived lineages from our cohort have the ability to enter cells expressing a low density of CD4, we performed SGA of full-length HIV-1 *env* genes using viral RNA as the template for cDNA synthesis followed by PCR done at template end-point dilution. Amplicons were sequenced to ensure that a single cDNA template initiated each amplification. A subset of the semen-derived HIV-1 *env* gene amplicons were cloned into an expression vector then used to pseudotype a virus made by cotransfecting with a Δ*env* HIV-1 backbone plasmid with a *renilla* luciferase reporter in order to produce pseudotyped virus that expressed participant-derived Env surface proteins. Pseudotyped virus was used to infect Affinofile cells that had been induced to express either high or low densities of CD4. The amount of luciferase produced by the cells was quantified and used as a surrogate measure of infectivity. As shown in Figure 5, semen-derived HIV-1 *env* genes, from both compartmentalized and equilibrated lineages, encoded Env proteins that require a high density of CD4 for efficient cell entry, indicating that they were being selected for replication in T cells.

**Figure 5.**
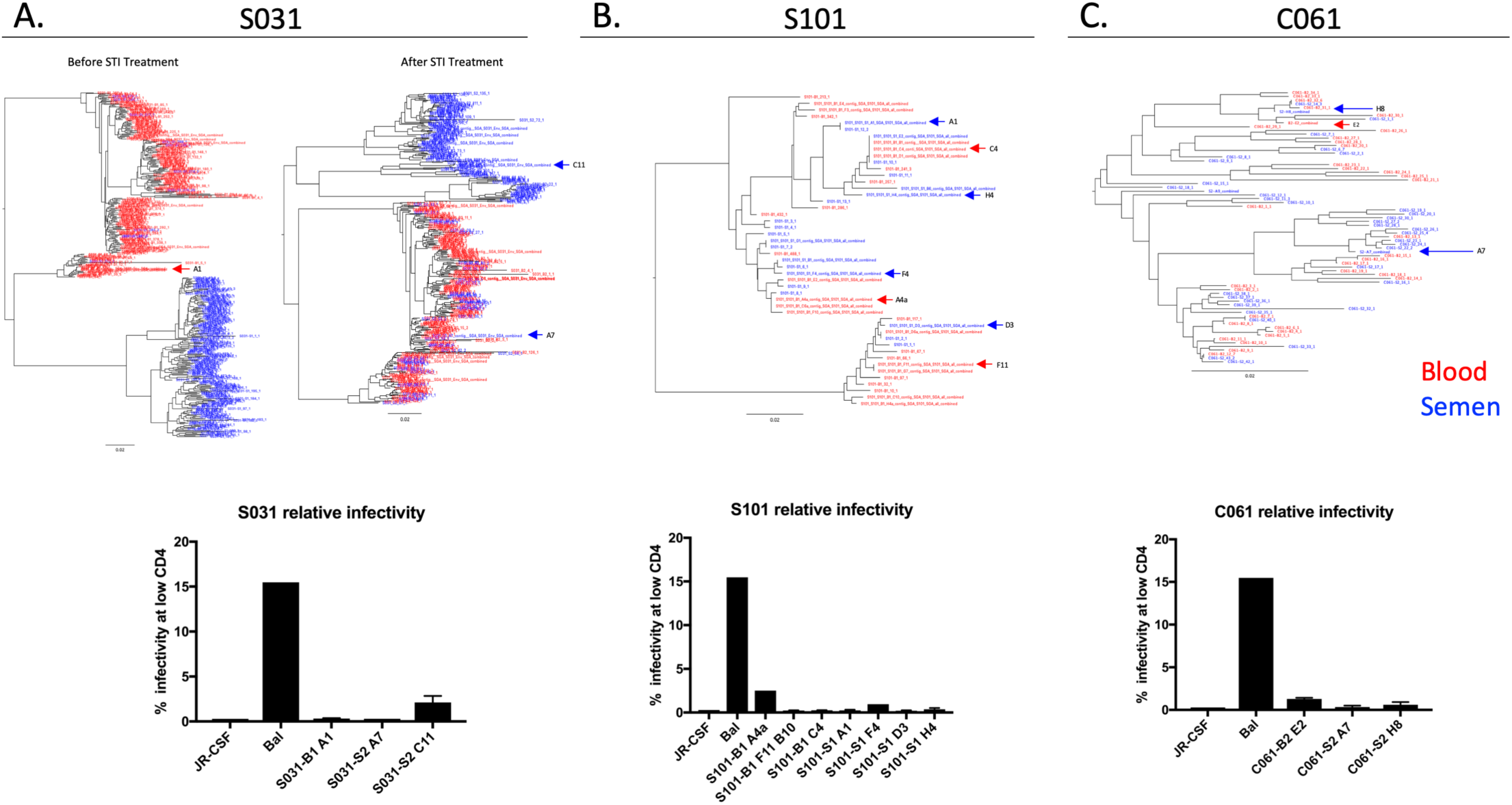
SGA-derived HIV-1 envelopes from the semen are T-cell tropic. (A-C) Neighbor-joining trees of *env* V1/V3 blood (red) and semen (blue) derived sequences. The graphs below depicts the ability of SGA-derived envelopes from blood and semen to enter cells expressing low densities of CD4. Colored arrows on the trees depict the locations of the envelopes used in the graphs below. JR-CSF and Bal are T-tropic and M-tropic controls, respectively. Data represent the average of three biological replicates.

### Cytokine/chemokine dynamics during treatment of the STI

To better understand the magnitude of the inflammation present within the genital tract during a concurrent sexually transmitted infection, we measured the concentrations of seven inflammatory cytokines and chemokines present in the blood and semen before and after treatment of the STI. To differentiate between STI-induced inflammation and HIV-induced inflammation, we included samples from HIV+ individuals not experiencing urethritis. As shown in Figure 6A, there was a group of cytokines (TNF-α, IL-6, and IL-1β) whose concentrations were increased in the semen of men with urethritis at the pre-treatment time point, and subsequently decreased after STI treatment. A second group of cytokines/chemokines, including CXCL10, IL-10, IFN-γ, and CCL2, were at similar concentrations in men with and without urethritis, as well as before and after STI treatment. A subset of four cytokines/chemokines (TNF-α, IL-10, CCL2 and CXCL10) were measured in blood as well (Figure 6B). There was no difference in the concentration of any of these analytes at any time point in men with or without urethritis, suggesting that STI-associated inflammation is limited to the genital tract and largely resolves with antibiotic treatment.

**Figure 6.**
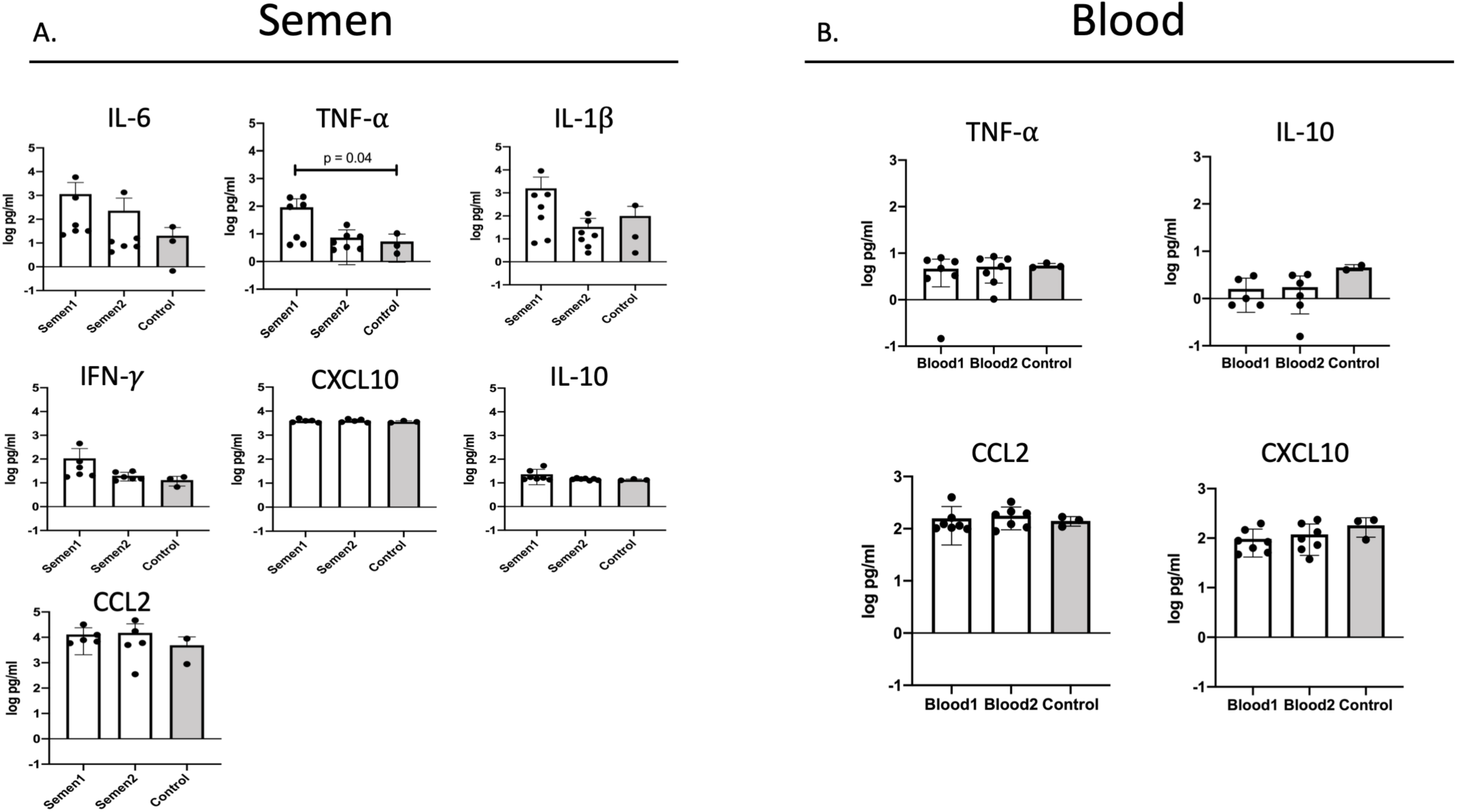
Cytokine/chemokine analysis in semen and blood before and after antibiotic treatment of the STI. Cytokine/chemokine concentrations in semen (A) and blood (B) were measured before and after STI treatment. The values were compared to a group of HIV+ men without a concurrent STI (labeled as “control”, and depicted by the grey bars). A one-way ANOVA was used to generate the indicated p value.

### Identification and characterization of a super-infection initiated in the male genital tract

In one participant with urethritis, S031, we observed a distinct, semen-only lineage in both the pre- and post-STI treatment time points (Figure 7A). Though the separate, semen-only lineage persisted across two time points, we did note that fewer semen-derived sequences were in semen-only clades in the second time point (post-STI treatment) as compared to the first time point (pre-STI treatment), suggesting that viral populations in the blood had been mixing with viral populations in the genital tract. Importantly, we observed semen-derived sequences that clustered with the blood-derived sequences at both time points, thus making contamination or sample mis-labeling an unlikely explanation for our observation (Figure 7A). When we constructed a neighbor-joining phylogenetic tree using blood- and semen-derived sequences from this participant and four others, we noted that the semen-derived sequences from S031 were as distinct from the blood-derived sequences as all participants were from one another, suggesting the presence of a superinfection (Figure 7B). A highlighter plot was used to identify the presence of recombinant lineages within the blood- and semen-derived sequences (Figure 7C), further supporting the notion of superinfection.

**Figure 7.**
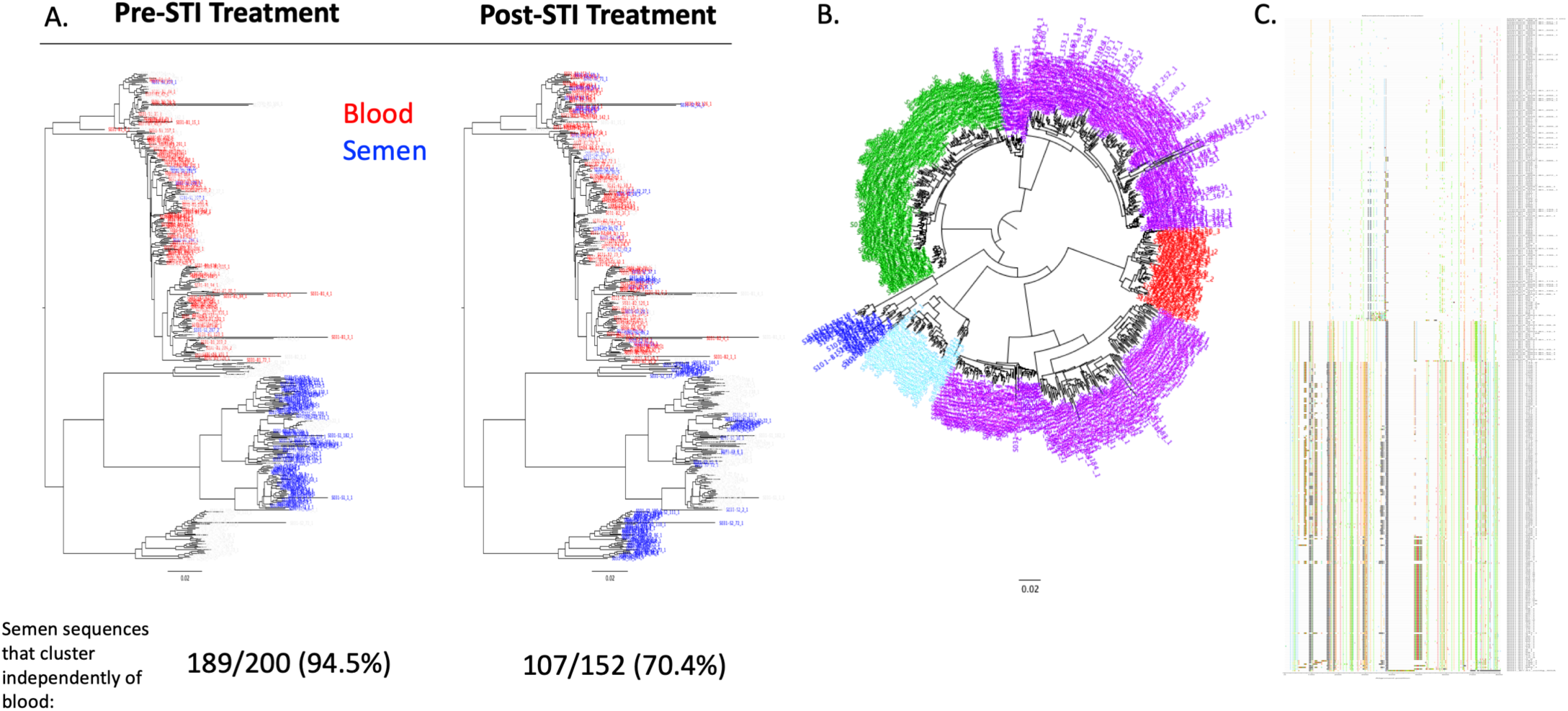
Characterization of a super-infection. A) Pre and post-STI treatment neighbor-joining phylogenetic trees depicting a distinct semen-only lineage from participant S031. Semen-derived sequences are shown in blue, blood-derived sequences are shown in red. The percent of all semen-derived sequences that cluster independently from the blood is shown at the bottom. B) A neighbor-joining phylogenetic tree containing blood and semen-derived sequences from five participants. Sequences from each participant are shown in a different color, sequences from S031 are purple. C) Highlighter plot depicting recombination between blood and semen-derived sequences from S031.

## Discussion

There have been numerous studies (12-14, 29-33) that have examined the prevalence of male genital tract compartmentalization of HIV-1, sometimes with discordant results. Some of these studies (11, 29, 30) examined the phenomenon of compartmentalization through the use of bulk amplification and/or cloning prior to sequencing; however these approaches have been shown to introduce sequencing artifacts, such as PCR-mediated recombination and sequence resampling (34–36). The use of deep sequencing with Primer ID in the current study corrects for PCR and sequencing errors through the creation of a template consensus sequence for each Primer ID-tagged cDNA (the template for PCR), while simultaneously allowing for the precise quantification of the total number of templates sequenced, i.e. the sample size of sampling of the viral sequence population (19). Thus, we can be confident that the viral variants we analyze are an accurate representation of the diversity found *in vivo*.

The observation that STI-associated urethritis does not significantly impact the degree of HIV-1 compartmentalization within the male genital tract raises several points. First, the mechanism underlying the establishment of compartmentalized lineages within the male genital tract remains unknown. This study sought to compare two possibilities: that STI-associated inflammation would lead to compartmentalized replication, or alternatively serve to recruit HIV-infected CD4+ T cells into the genital tract, thereby equilibrating the viral populations found in the blood and semen. We observed compartmentalization in 26% of men with urethritis, and 20% of men without urethritis; thus, given this number of participants we did not detect a difference in the extent of compartmentalization with and without and STI-associated urethritis. In the overall cohort, we observed compartmentalization in the genital tract of 25% of men. This prevalence of compartmentalized lineages in the genital tract is similar to what was observed in a previous study (9) that used a heteroduplex tracking assay to examine the relationship between blood and semen-derived *env* V3 populations in men with and without urethritis. In this earlier study, they observed discordant V3 populations between the blood and semen of 40% of men. Importantly, there was no difference in the V3 population dynamics between the blood and semen of men with urethritis, compared to men without urethritis. Later, Anderson and colleagues (10) utilized single genome amplification to examine the relationship between blood and semen-derived HIV-1 envelopes in men without urethritis. Here, they reported a 31% prevalence of compartmentalization in the genital tract. They also observed clonal amplification in the semen of men without urethritis. Compartmentalized populations in the genital tract have also been observed in the context of acute HIV-1 infection. In a recent study by Chaillon *et al.* (31), deep sequencing was used to examine HIV-1 populations in blood and semen in early infection. They observed compartmentalization in 2 of 6 participants at baseline (a median of 81 days after the estimated date of infection).

The second noteworthy point pertains to the source of HIV-1 shed in the semen. HIV-1 (12, 13, 37) and/or SIV (38) RNA has been recovered from a variety of male genital tract tissues including the urethra, prostate, testis, seminal vesicles, vas deferens and epididymis. Our observation that inflammation does not alter the frequency with which we detect semen-specific HIV-1 lineages suggests that, when compartmentalized lineages are present, they are most likely produced by cells in anatomical areas that are not in direct contact with the periphery. In one extreme case of this type of isolation we previously observed the presence of a macrophage-tropic variant in semen (28) which suggests that in this case there was sufficient depletion of CD4+ T cells, without replenishment, that the virus evolved to expand its target cell specificity. In the current study all of the viruses tested were T cell-tropic, requiring a high density of CD4 for efficient entry into cells. In addition, all were predicted to use CCR5 as a coreceptor based on genotypic predictions of the V3 loop sequence (data not shown). This result is important as a recent report by Ganor and colleagues (39) reported the presence of macrophage-tropic viral variants in urethral tissues, suggesting the possibility of a urethral reservoir. However, it appears such variants are not shed in the semen.

Compartmentalization in the male (30, 31) genital tract has largely been defined as a transient phenomenon. Here, we examined how antibiotic treatment of a concurrent sexually transmitted infection (primarily gonorrhea or trichomonas) impacted the relationship between blood- and semen-derived HIV-1 *env* V1/V3 sequences. We found that viral variants present before STI treatment remained detectable after STI treatment, and furthermore, that the relationship between blood and semen-derived sequences remained consistent throughout the course of STI co-infection. In only one participant out of 13 did we detect a change in the relationship between blood and semen-derived sequences over time. In this instance, the depth of sampling pre-STI treatment was relatively poor, with only 13 V1/V3 sequences recovered per compartment, while the sampling post-STI treatment was much greater (103 sequences per compartment). Thus, it is quite possible that the relatively few sequences obtained pre-treatment obscured the presence of the compartmentalized lineage that we observed post-treatment. It is also important to note that while gonococcal infections are cleared rapidly from the urogenital tract after a single antibiotic treatment (40), the underlying immune activation can persist, as demonstrated by the fact that in men with an STI, HIV-1 viral loads in semen were still higher than in men without an STI, even after effective antibiotic treatment (20), although we were able to measure some diminution of inflammation with a change in some inflammatory markers. Therefore, while we do observe stable relationships between blood and semen-derived sequences both before and after STI treatment, our conclusions are limited by the relatively short period of follow-up. It is worth noting that in one participant, virus in the semen was compartmentalized relative to the blood both before and after STI treatment but the compartmentalized lineage in the semen changed between the two time points. Both lineages, while minor, were complex in sequence composition and thus the latter one did not evolve over the short period of time between the two samplings. Thus, there must have been reduced production of one lineage and the appearance of a pre-existing lineage over a relatively short period of time.

Given the error-prone nature of HIV-1 reverse transcription (41), a single transmitted/founder viral variant rapidly evolves into a diverse population within an infected individual (42–44). As such, the identification of identical or nearly identical sequences in the blood in chronic untreated infection is relatively infrequent. There are two mechanisms to consider that can explain the presence of such sequences. In people on therapy there can be low level production of virus particles with identical sequences and this is thought to be due to clonal expansion of an infected cell (45–47) some of which can produce a low level of infectious virus (48, 49). In the absence of therapy, the viral load of viruses with similar sequences is much higher, suggesting either that the corresponding cellular expansion is much greater or that the virus comes from another source, i.e. replication, after passing through a recent genetic bottleneck. This question becomes even more relevant using the shorter amplicon associated with deep sequencing as a significant fraction of the viral sequences cluster into lineages of identical sequences. In order to determine if the identical sequences from deep sequencing observed off therapy in these men were truly clonal, we compared sequences obtained from deep sequencing to those obtained as full length *env* genes using template end-point dilution PCR (SGA). We found that the sequences that were identical in the deep sequencing data set were in a population of similar but not identical sequences when the larger region of the genome was analyzed (Figure 4, Supplemental Figure 1-3). We conclude that these populations are present at their detected level due to ongoing viral replication. However, the high level of similarity in these sequences implies a recent genetic bottleneck prior to expansion by viral replication, although the nature of that bottleneck remains unknown and could still be due to clonal expansion of an infected cell amplified by a burst of local replication. It is possible that this phenomenon is mediated by an infected antigen-specific cell that undergoes amplification due to the presence of the STI.

HIV-1 infection is associated with dysregulation of seminal cytokines (50, 51) as well as an increased semen: blood cytokine ratio (10, 50, 51). This pro-inflammatory environment has been suggested to increase viral replication as semen viral load often correlates with cytokine levels (52), as well as the fact that several cytokines, including TNF-α, directly act on the virus to increase replication (53) (reviewed in (54)). A similar phenomenon is observed in men with classical STIs such as gonorrhea or trichomonas (20). We analyzed cytokine/chemokine levels in the blood and semen of men with and without STI-associated urethritis to determine if inflammation increased with the presence of a concurrent STI infection and whether such inflammation had resolved during the two-week period of follow-up. Among the seven cytokines/chemokines analyzed (IL-6, TNF-α, IL-1β, IFN-γ, CXCL10, IL-10 and CCL2) only TNF-α was significantly increased in the semen of men with STI-associated urethritis, compared to HIV-positive men without urethritis. However, levels of IL-6 and IL-1β were also increased in men with urethritis, though the difference was not statistically significant. Importantly, the levels of TNF-α, IL-6 and IL-1β all decreased to levels similar to that of men without urethritis after STI treatment. Thus, we observed that men with urethritis have an enhanced pro-inflammatory environment compared to HIV-positive men without urethritis, and that this difference is reduced following antibiotic treatment of the STI. As expected, cytokine levels in the blood were similar in men with and without STI-associated urethritis and remained unchanged following STI treatment. This result further supports our finding that inflammation due to STI-associated urethritis does not impact the formation of compartmentalized lineages in the male genital tract.

## Acknowledgements

We would like to acknowledge all participants. We would also like to thank Li-Hua Ping for her help in organizing the samples and data. This work was supported by NIH award 5R01DK108424-05 to MSC, and by R01AI140970 to RS. ODC was supported by NIH training grant T32-AI007001. The work was also supported by the UNC Center For AIDS Research (NIH award P30 AI50410) and the UNC Lineberger Comprehensive Cancer Center (NIH award P30 CA16068). We wish to acknowledge the efforts of the UNC High Throughput Sequencing Facility.

